# Histone deacetylase activity is required for *Botrylloides leachii* whole body regeneration

**DOI:** 10.1101/433854

**Authors:** Lisa Zondag, Rebecca Clarke, Megan J Wilson

**Affiliations:** Developmental Biology and Genomics Laboratory, Department of Anatomy, Otago School of Medical Sciences, University of Otago, P.O. Box 56, Dunedin 9054, New Zealand

**Keywords:** Regeneration, epigenetics, tunicate, histone

## Abstract

The colonial tunicate *Botrylloides leachii* is exceptional at regenerating from a piece of vascular tunic after loss of all adults from the colony. Previous transcriptome analyses indicate a brief period of healing before regeneration of a new adult (zooid) in as little as 8–10 days. However, there is little understanding of how the resulting changes to gene expression, required to drive regeneration, are initiated and how the overall process is regulated. Rapid changes to gene expression often occur in response to chromatin changes, mediated by histone modifications such as histone acetylation. Here, we investigated a group of key epigenetic modifiers, histone deacetylases (HDAC) that are known to play an important role in many biological processes such as development, healing and regeneration.

Through our transcriptome data, we identified and quantified the expression levels of HDAC and histone acetyltransferase (HAT) enzymes during whole body regeneration (WBR). To determine if HDAC activity is required for WBR, we inhibited its action using valproic acid (VPA) and Trichostatin A (TSA). HDAC inhibition prevented the final morphological changes normally associated with WBR and resulted in aberrant gene expression. *B. leachii* genes including *Slit2, TGF-*□*, Piwi* and *Fzd4* all showed altered gene expression upon HDAC inhibition in comparison to the control samples. Additionally, atypical expression of *Bl_Piwi* was found in immunocytes upon HDAC inhibition.

Together, these results show that HDAC function, specifically HDAC I/IIa class enzymes, are vital for *B. leachii* to undergo WBR successfully.

## Background

*Botrylloides leachii* whole body regeneration (WBR) requires a series of rapid molecular and cellular responses as a consequence of loss of all adults (termed zooids). Regeneration of a new zooid can occur in as little as 8–10 days from a small vascular fragment (Rinkevich et al., 1995). RNA-sequencing (RNA-seq) followed by differential gene expression analysis between early and late regeneration stages revealed novel information on gene expression changes during WBR (Zondag et al., 2016). The expression of genes with known roles in wound healing, cellular organisation, and developmental pathways such as transforming growth factor □ (TGF-□) and Notch signalling, all changed significantly over the 8 day WBR period (Zondag et al., 2016).

We predict that the immediate changes to gene expression required for *B. leachii* WBR requires an epigenetic mechanism. Modifications to histone proteins, which make up the nucleosome complex, along with chemical additions directly to the DNA such as methylation at CpG sites, represent key mechanisms of gene regulation (Gan et al., 2007; Jopling et al., 2011). These epigenetic modifications function by inducing changes in chromatin structure, either by permitting or restricting DNA accessibility or through the recruitment of DNA binding proteins (Bannister and Kouzarides, 2011).

The chemical modification of histone proteins is a dynamic process requiring enzymes to add and remove ‘marks’ to specific amino acids at the N-terminus of histone proteins. Two well-studied proteins that regulate acetylation of histone proteins are histone acetyltransferases (HAT) and histone deacetylases (HDAC). Different classes of HAT enzymes transfer an acetyl group onto specific lysine residue located within the histone(s) protein target (Lee and Workman, 2007). Generally, HATs promote transcriptional activation through the creation of binding sites for chromatin remodelling complexes to bind and open up the chromatin (Marmorstein and Roth, 2001). HDACs are thought to oppose HAT activity, repressing transcription by removing acetyl groups from histone tails which causes the chromatin to move towards a heterochromatic state (Gregoretti et al., 2004; Murakami, 2013). Condensing of the DNA around tightly packed nucleosomes prevents transcription factors from accessing transcriptional start sites and therefore represses local gene expression (Ke et al., 2012). By controlling the acetylation state of histones, HATs and HDACs are involved in regulating transcription of genes and consequently the regulation of many processes including cell proliferation, apoptosis, tumorigenesis, and cell differentiation (Ke et al., 2012).

Cellular reprogramming and re-establishment of lost cell types is a critical process during regeneration, and consequently HDAC activity is required for successful regeneration in some vertebrates (Huang et al., 2013; Taylor and Beck, 2012; Tseng et al., 2011) and planarians (Eisenhoffer et al., 2008; Reddien et al., 2005; Robb and Sanchez Alvarado, 2014). HDAC inhibition (HDACi) during *Xenopus* tail regeneration by Trichostatin A (TSA) and valproic acid (VPA) halts regeneration (Taylor and Beck, 2012; Tseng et al., 2011). Ectopic expression of *BMP2* and *Notch1* genes, two genes required for *Xenopus* tail regeneration, was also observed following HDACi (Tseng et al., 2011). In mammalian models, VPA treatment also inhibits the regenerative response; for example, mice exposed to HDAC inhibitors are impaired in their ability to regenerate their liver following resection (Huang et al., 2013; Ke et al., 2012). Here, HDACi prevented hepatocyte cell proliferation, and altered expression of cell cycle genes, for example resulting in an upregulation of *b-myc*, an inhibitor of cell proliferation, and decreased expression of *cyclin D1*, suggesting an essential role for HDAC in regulation of gene required for cell cycle progression (Ke et al., 2012). Further investigations found that changes to global acetylation following HDACi paralleled changes in gene expression, with some genes normally induced upon initiation of liver regeneration showing reduced expression with HDACi, and genes normally suppressed during liver regeneration having increased expression following VPA treatment (Huang et al., 2014).

In contrast, little research to date has been carried out analysing the specific role of HAT proteins in regards to regeneration. It has been shown that p300 (a HAT belonging to KAT3 family) when overexpressed allowed axonal regeneration to occur after optical nerve damage in rats (Gaub et al., 2011), likely through the reactivation of genes important for regeneration. Shubuya *et al*. (2015) demonstrated that regeneration was still possible in the tunicate species Stolidobranchian, *Polyandrocarpa misakiensis*, when HAT (GCN5) was inhibited with CPTH2 (cyclopentylidene [4'-(4-chlorophenyl)thiazol-2-yl) hydrazone). This suggests that *P. misakiensis* either has a built-in redundancy for the targeted HAT protein or that it is not essential for regeneration. However, CPTH2 exposure during budding,a mechanism of asexual reproduction in these species, caused down-regulation of trans-differentiation-related genes (Shibuya et al., 2015), implying that HAT is important for asexual reproduction.

Regenerative studies looking at HDAC and other epigenetic modifiers demonstrate their importance in allowing appropriate reestablishment of the tissue by up-regulating both cell proliferation and growth, followed by appropriate cell differentiation. Any alterations to the activity or expression of these chromatin modifiers could cause disruption to the acetylation homeostasis (Haberland et al., 2009; Saha and Pahan, 2006) and trigger aberrant gene expression. Therefore, epigenetic processes are of interest to studying regenerative mechanisms in many species, but this has been largely unexplored in tunicate models such as *B. leachii*.

The current hypothesis is that the cells lining the vascular vessels in a *B. leachii* colony act as the ‘stem’ cell population for WBR (Rinkevich et al., 2010). To allow the putative ‘stem’ cells (attached to vascular lining) in *B. leachii* to become activated and enter a proliferative state at the onset of WBR, the chromatin architecture likely needs to be modified to allow a rapid shift in gene expression to occur. Once activated these cells move into the vascular lumen and are predicted to be the progenitor source of many of the different cell types of the new zooid during WBR (Rinkevich et al., 2010). We hypothesize that during WBR cell fate activation, differentiation and/or reprogramming of somatic cells occurs through chromatin modifications. To test this, we firstly identified *HDAC* and *HAT* genes in the *B. leachii* reference transcriptome (Zondag et al., 2016), and analysed their expression across regeneration stages, using transcriptome and RT-qPCR analysis. We found that *HDACI* and *HDACII* genes were expressed at high levels throughout WBR. Secondly, inhibition of HDACI/II activity either by VPA or TSA halted the WBR process and resulted in changes to the expression of key regeneration-related genes including *Piwi*, a gene used as a marker of stemness.

## Results

### Identification and classification of *B. leachii HDAC* and *HAT* genes

To initially determine if HDAC and HAT enzymes are expressed during WBR, we identified transcripts for candidate orthologues within the regeneration transcriptome (Zondag et al., 2016) and further confirmed their identities using phylogenetics. For phylogenetic analysis we also included protein sequences from additional ascidians with sequenced genomes, one representing a solidary ascidian *Ciona robusta* (formerly known as *C. intestinalis* type A (Gissi et al., 2017)), and *Halocynthia roretzi, Molgula oculata* and *Botryllus schlosseri*, representing additional Stiolidobranchians. Candidate sequences from the *B. leachii* regeneration transcriptome and the *B. schlosseri* genome were identified by tBLASTn using conserved HAT and HDAC protein domains (File S1) and confirmed by reciprocal SMARTBLAST.

There are four main classes of HDACs. The first class, Class I, contains HDAC1, 2, 3 and 8. Class II contains HDAC 4, 5, 6, 7, 9 and 10, Class III includes the unrelated Sirtuin (SIRT) family (NAD-dependent enzymes) and Class IV contains HDAC 11 ((Gregoretti et al., 2004), Fig. 1 and Supplementary Table 1). The classical HDAC proteins are considered to have descended from a common ancestor (Gregoretti et al., 2004). We identified a total of six candidate transcripts expressed during WBR that encode HDAC-type proteins (Supplementary Table 1). Phylogenetic analysis clustered these into separate clades, with representative sequences from mouse, human, and ascidian genomes. We identified three Class I type HDAC proteins (named Bl_HDAC3, Bl_HDAC8 and Bl_HDAC2) within the *B. leachii* regeneration transcriptome. All three are also present in *C. robusta, H. roretzi* and *M. oculata* genomes, and two within the *B. schlosseri* genome (Bs_HDAC2 and Bs_HDAC8; Fig. 1). The class I HDAC proteins clustered tightly together, supporting a high level of sequence conservation for these proteins.

**Figure 1:**
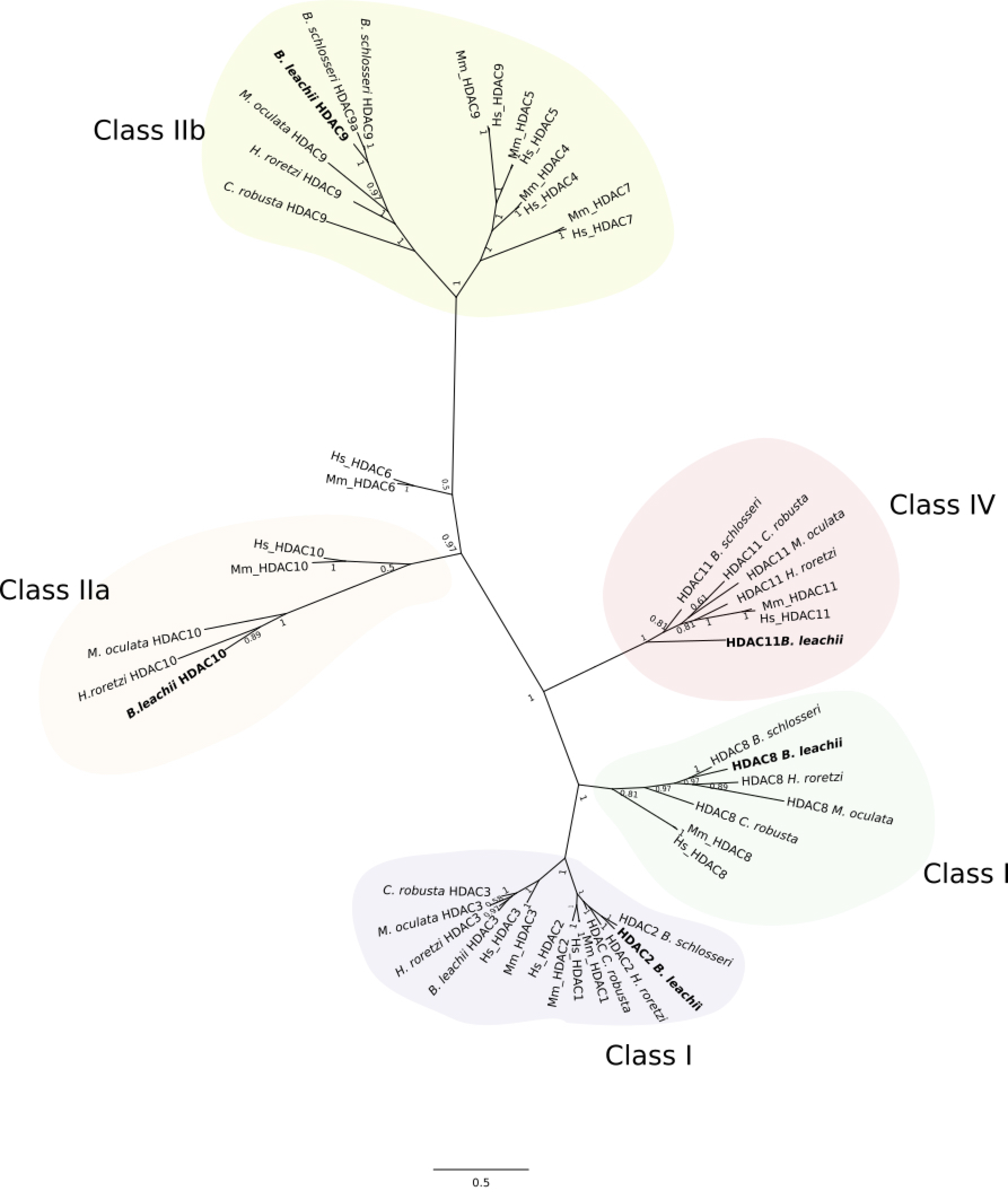
Classification of *B. leachii* HDAC and HAT genes. Bayesian phylogeny for HDAC proteins displayed with FigTree (Ronquist et al., 2012). Sequences identified from *Botrylloides leachii* regeneration transcriptome, along with additional HDAC sequences from *B. schlosseri, M. oculata, H. roretzi* and *C. robusta* were used to construct the molecular phylogenetic tree to classify *B. leachii* HDAC proteins. Class I, II and IV all refer to the different HDAC classes. Abbreviations: *Homo sapiens* (*Hs*)*, M. musculus* (*Mm*). Node labels are posterior probabilities.

Class II proteins, HDAC4/5/7/9 and HDAC6/10 are subclassified into Class IIa and ClassIIb respectively (Fig. 1; (Gregoretti et al., 2004)). Two *B. leachii* HDAC proteins were classified as Class II HDACs: HDAC9 (IIa) and HDAC10 (IIb; Fig. 1). HDAC10 and HDAC5/9 proteins formed distinct groupings, suggesting more sequence divergence between class II HDAC proteins. HDAC5, 4, 7 and 6 are found only in the vertebrates, and are thought to have arisen by gene duplication after divergence of vertebrates and invertebrates (Gregoretti et al., 2004). Supporting this hypothesis, only a single copy of each Class IIa and IIb were found in ascidian genomes, with the exception of *B. schlosseri*, which has a duplication of HDAC9 (Fig. 1). Lastly, a single Class IV protein, HDAC11 was identified in *B. leachii* and other ascidian genomes (Fig. 1).

HAT/KAT proteins are divided into subgroups through differences in sequence homology and function (Liew et al., 2013). Major HAT protein families include the MYST, GCN5-related proteins (GNAT) and the p300 family (Sapountzi and Cote, 2011). A total of seven HAT-related transcripts were identified in a BLAST search of the *B. leachii* transcriptome (Fig. 2 and Supplementary Table 1). As each HAT protein can have distinct cellular functions we carried out phylogenetic analysis to classify into subgroups, along with vertebrate and four closely related ascidians to *B. leachii*.

**Figure 2:**
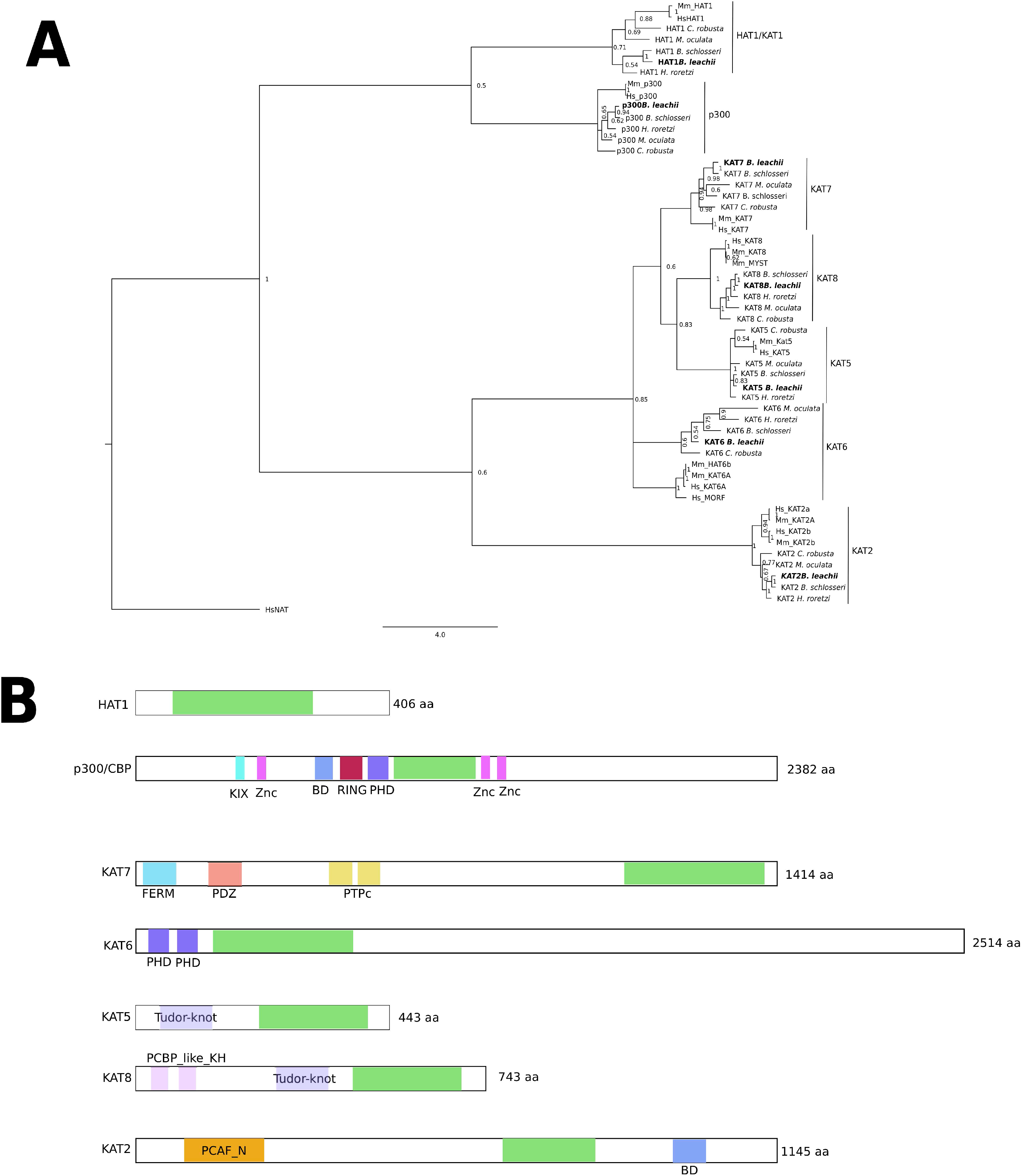
Classification of *B. leachii* HAT/KAT genes. Identified *B. leachii* HAT/KAT genes were classified through phylogeny and the identification of additional protein domains. **A**. Bayesian phylogeny for HAT proteins displayed with FigTree (Ronquist et al., 2012) with additional HAT sequences from *B. schlosseri, M. oculata, H. roretzi* and *C. robusta*. Hs_NAT (N-acetyltransferase) was used to root the tree. Node labels are posterior probabilities. **B**. Protein domains present in *B. leachii* KAT proteins. The acetyltransferase catalytic domain (green) is flanked by additional protein domains important for conferring functional specificity to each KAT/HAT protein. Abbreviations: *Homo sapiens* (*Hs*)*, M. musculus* (*Mm*). Bromodomain (BD), PHD (Plant HomeoDomain) Znc (Zinc binding domain), FERM (Four-point-one, Ezrin, Eadixin, Mosesin), RING (Really Interesting New Gene), PCBP_like_KH (PolyC Binding Protein hnRNP K Homology), PCAF_N (p300/CBP-associated Factor N-terminal)

Phylogeny analysis showed that within the *B. leachii* WBR transcriptome, orthologues for five MYST genes, *KAT2, KAT5, KAT6, KAT7, KAT8*, a single *p300* gene (*Bl_p300*) and a single GNAT (*Bl_HAT1*), are present. The KAT/HAT proteins separate into 7 distinct groups with a single representative from each ascidian genome (Fig. 2A). Within most clades, the ascidian sequences clustered separately from the vertebrate group suggesting that they are more closely related (Fig. 2A). In addition to the catalytic domain, HAT/KAT proteins contain further protein domains important for mediating molecular functions and impart some specificity in KAT recruitment to particular chromatin sites, often termed “reader” domains (Yun et al., 2011). As these additional protein domains are important for function, we identified KAT reader domains for *B. leachii* proteins (Fig. 2B, Supplementary Table 3). Bl_HAT1 is the smallest KAT protein with no identifiable addition reader domains. HAT1 is often referred to a Type B histone acetyltransferase; enzymes that acetylate free histone proteins before assembly into nucleosomes, with Type A histone acetyltransferases associated with chromatin binding (Parthun, 2007). Both Bl_p300 and Bl_KAT2 both contain bromodomains, this region is important for the assembly of protein complexes and recognition of acetyl lysine (Dhalluin et al., 1999). Bl_KAT8 and Bl_KAT5 have tudor-knot domains, these mediate chromodomain interactions with RNA (Shimojo et al., 2008). We predict that Bl_KAT7 is likely to be a cytoplasmic protein due to the presence of the FERM domain ((Chishti et al., 1998), Fig. 2A).

### *B. leachii HDAC* and *HAT* genes are consistently expressed during WBR

We next analysed the expression levels of each *HAT* and *HDAC* genes during WBR both in our previous RNA-seq data sets (File S2 (Zondag et al., 2016)), and by RT-qPCR for highly expressed transcripts that are of further interest (Fig. 3).

**Figure 3:**
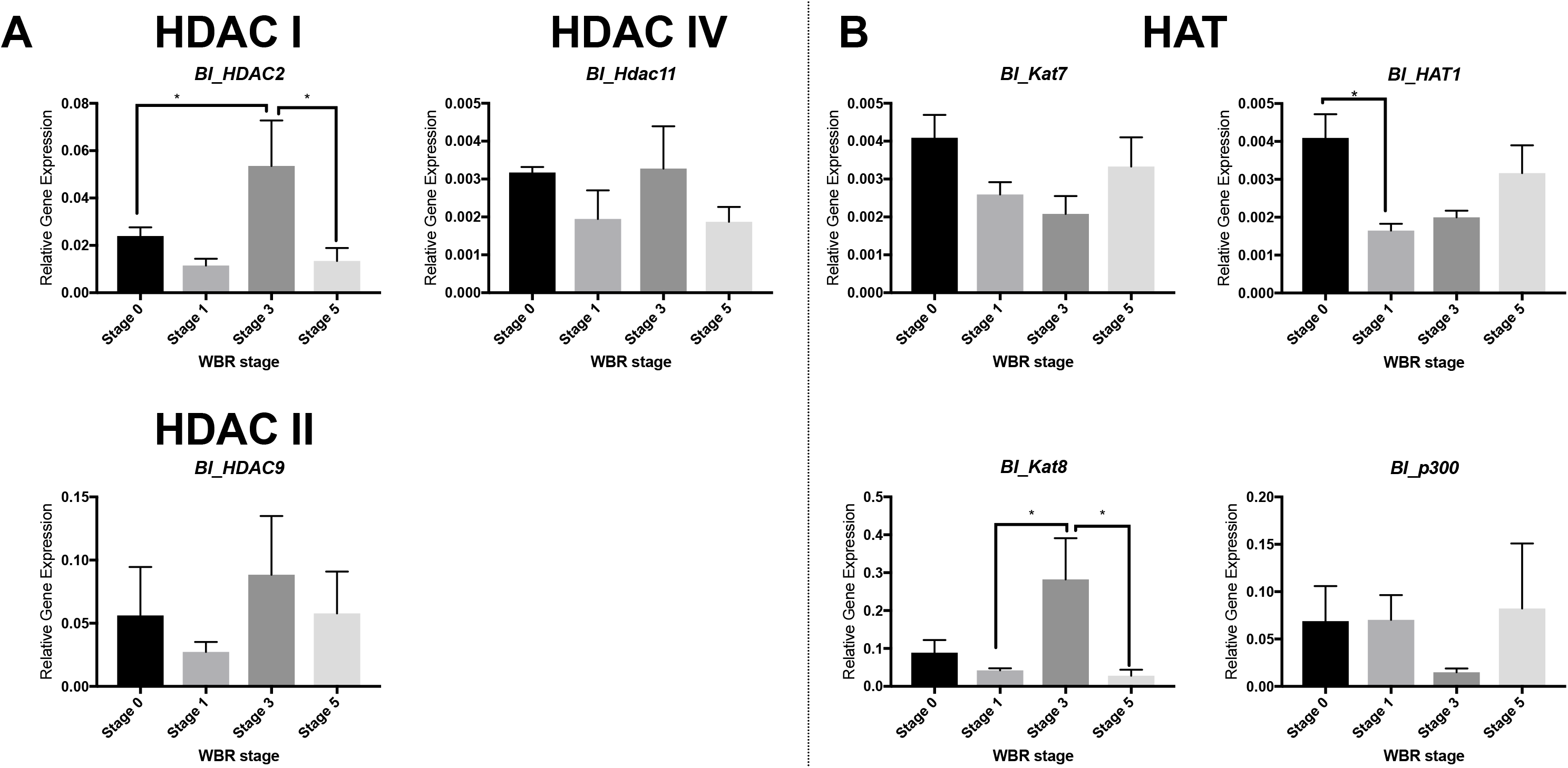
*B. leachii HAT* and *HDAC* gene expression during WBR. Relative expression levels for *HDAC* (A) and *HAT* (B) transcripts across three WBR stages following zooid removal. **A**. WBR expression shown for HDAC IV-type gene, *Bl_HDAC11;* HDAC II class gene *Bl_HDAC2;* and HDAC III class gene *Bl_HDAC9*. **B**. Expression levels for *HAT* genes: *Bl_p300, Bl_HAT1, Bl_Kat8* and *Bl_Kat7*. Data shown as mean +/− standard error of the mean. Statistical analysis was carried out using a one-way ANOVA, with Tukey’s multiple comparison correction (P < 0.05 = *).

HDAC transcripts showed some variation in expression levels across WBR and even between biological replicates (Fig. 3A). *B1-HDAC2* (HDAC class I) was expressed at higher levels than the other *HDAC* class genes (File S2; > 100 FPKM). Expression of *B1-HDAC2* significantly increased between stages 0 and 3 (P = 0.02, Padj = 0.01, one-way ANOVA, Tukey’s multiple comparisons test), before declining again just prior to the appearance of the new zooid between stages 3 and 5 (Fig. 3A; P = 0.04). *B1-HDAC9* gene expression remained at a similar level across all examined timepoints (Fig. 3A). The *B. leachii* HDAC class IV gene (*B1*_HDAC11) was expressed at similar levels across all regeneration stages (Fig. 3A).

All seven *HAT* genes were expressed at varying levels during WBR (File S2; Fig. 3B), we confirmed expression for the top four genes by RT-qPCR. *Bl_p300* transcript expression remained at similar levels through the first four stages of WBR, before a slight increase between stages 4 and 5 (~96 - 168 h; Fig. 3B). *Bl_HAT1* expression declined during the first 15 h of WBR (Fig. 3B). *B1-KAT7* gene expression was maintained at a similar expression level across WBR (Fig. 3B and File S2). *Bl_KAT8* expression increases between stages 1 and 3, to peak at stage 3 (P = 0.02; Fig. 3B) before declining back to expression levels similar to earlier stages at stage 5 (P = 0.04; Fig. 3B). *Bl_KAT6, Bl_HAT2A* and *Bl_KAT5* were expressed at much low levels throughout WBR (File S2, < 10 FPKM).

### HDAC inhibition halts *B. leachii* whole body regeneration

HDAC enzymes of class I and II had the highest level of gene expression in the HDAC family (Fig. 3A). To determine if HDACI/II function is required for successful WBR, we used an HDAC chemical inhibitor. VPA is an inexpensive and established inhibitor of HDAC class I/IIa (Gottlicher et al., 2001). It is thought to block HDAC activity by displacing the zinc ion interacting with the catalytic site of HDAC, thereby prevent it from carrying out histone de-acetylation (de Ruijter et al., 2003). As such, successful HDAC inhibition would be expected to increase acetylation levels of cellular proteins. We confirmed that VPA treatment increased total acetylation levels for nuclear proteins using an antibody raised against acetyl-peptides (anti-Pan-acetyl). The total amount of acetylated protein increased by ~2.5 fold (Fig. S2).

While new zooids were normally observed by day 10, HDAC inhibitor experiments were performed for a total of 18 days to allow a definite conclusion to the stage of WBR reached by the vascular fragments in the VPA treated and control groups reached. The majority (69%) of VPA treated vascular fragments died during stages 3 and 4 (Fig. 4A). Only 1 of the regenerating fragments developed past stage 4 (siphons in the developing zooid had started to emerge). However, malformations were seen at stage 4 as the regeneration niche was merging with surrounding tissue (Fig. 3B) and between stages 4 and 5, where abnormal siphons and darkening of the fragment indicated it was not healthy, and this fragment subsequently died (Fig. 4B).

**Figure 4:**
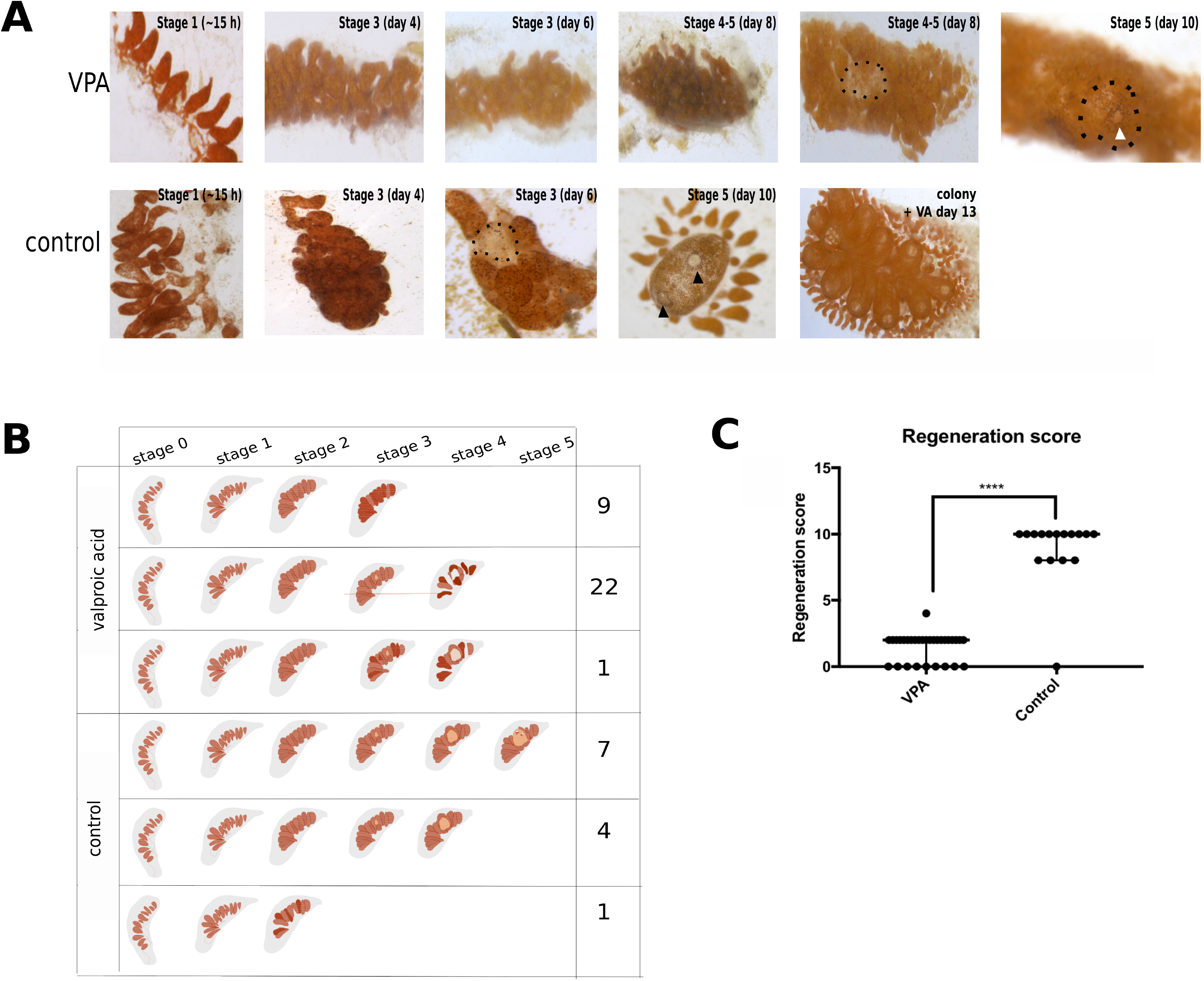
HDAC inhibition prevents completion of WBR. *B. leachii* vascular tissue exposed to VPA showed successful progression of WBR until stage 3. **A**. Example images of regenerating tunicate fragments in the presence of VPA. Most fragments showed signs of death (darkening and covered in a white film) around days 4–6. One *B. leachii* vascular fragment exposed to VPA reached stage 4, as a regeneration niche is visible (arrow), however, a malformed siphon started to develop (stage 5, white arrowhead) before the regenerating tissue starts darkened and died. Control fragments showed a typical WBR process, with a regeneration niche (dashed line) forming around ~days 5–6 and a zooid at day 10 (black arrowheads indicate siphons). Intact colonies exposed to VPA for the same length of time were healthy. **B**. Summary of the stages during WBR that vascular fragments exposed to VPA (or control) reached after 18 days of culturing. **C**. Regeneration scores of HDACi experiments after leaving the vascular fragments for a total of 18 days. There was a significant difference (Mann-Whitney test P value <0.0001) between regeneration score results of the VPA exposed fragments versus the controls. Data shows median ± interquartile range. P≤ 0.0001 (****), P ≤ 0.001 (***), P ≤ 0.01 (**), P ≤ 0.05 (*), P > 0.05 (ns)

To further examine the effect of HDAC inhibition with VPA, regenerating fragments were collected at 48 h and 5 days post zooid removal for histology (Fig. 5). Typically, by 48 h regenerating vascular fragments have reached stage 3, and regeneration niches have formed often containing a regenerating bud (Fig. 5Ai-iii). However, in VPA-treated fragments, areas where ampullae have fused either had a bud that appeared abnormal (Fig. 5vi and vii), or did not contain evidence of a regeneration bud anywhere in the vascular fragment (Fig. 5A iv and v). Within control regenerating fragments left for 5 days, the regenerating bud zooid had formed the rudiments of branchial chamber and gut chambers (Fig. 5B, i and ii). Whereas, VPA-treated fragments collected at the same time-point that did form a regeneration bud failed to further develop, and these buds appeared to be either degenerating or had not formed at one end (Fig. 5B iii and iv asterisk).

**Figure 5.**
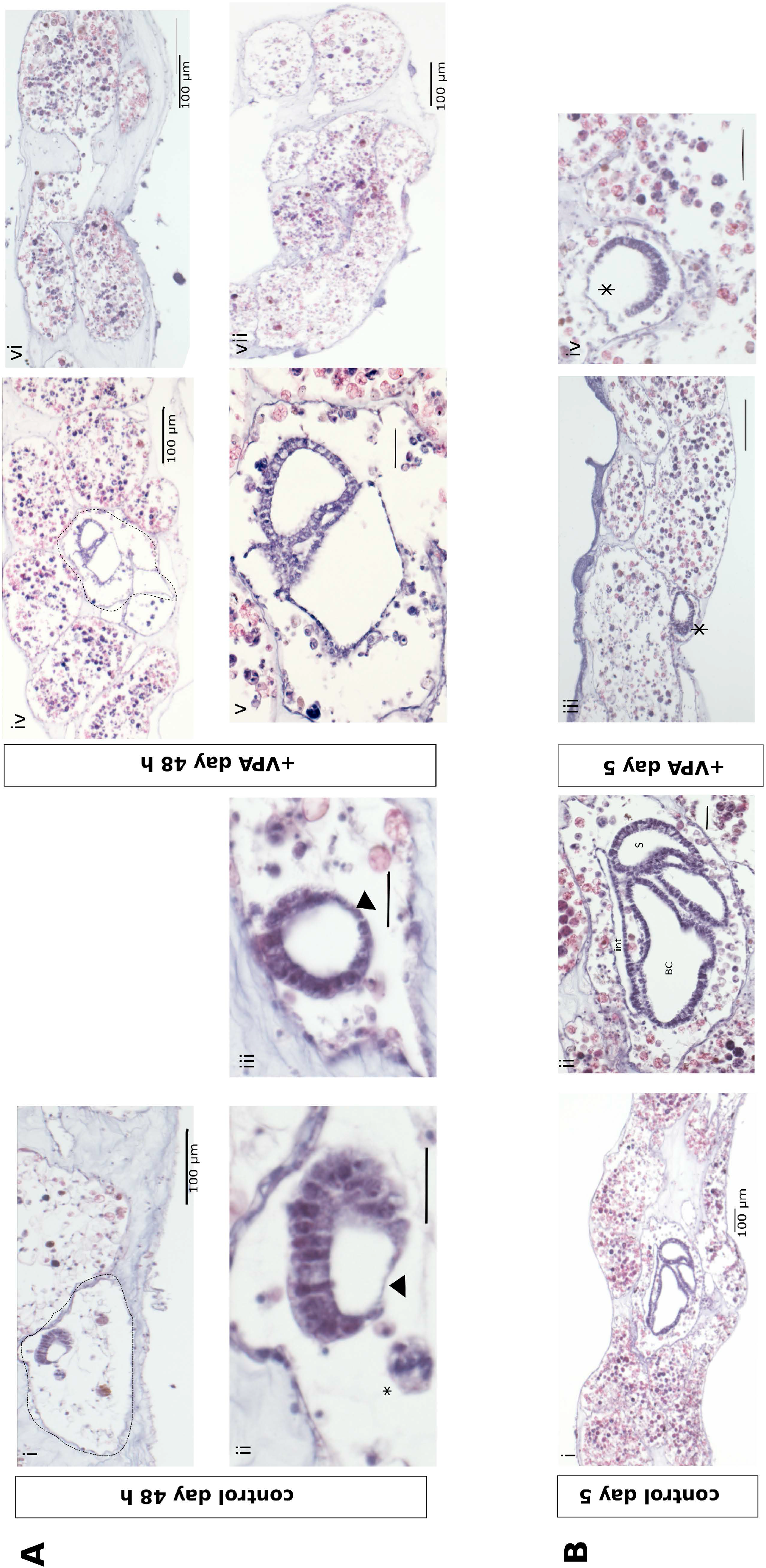
Histology of control and VPA-treated fragments. **A**. Control (i-iii) and VPA-treated (iv-vii) fragments collected at 48 h. At this stage of regeneration, control fragments have formed regeneration niches (dashed line, i and iv), containing a epithelial bud that is thinning on one side (ii and iii, arrowhead). Often a cell aggregate (ii, asterisk) is observed nearby (depending upon the level of section through the fragment). Most VPA-treated fragments fail to form a regeneration bud at 48 h (vi and vii), and those that contain a bud appear to be malformed (iv and v (higher magnification)). **B**. By day 5, control fragments have a regenerating zooid, with rudiment tissue layers forming (i and ii), whereas, VPA-treated fragments either contain no buds or a degenerating malformed bud (iii and iv, asterisk). Scale bars indicate 25 μm unless otherwise indicated. Abbreviations: branchial chamber (BC), stomach (S) and intestine (int).

Finally, to additionally confirm that HDAC activity was needed for successful regeneration, we used a second known HDAC inhibitor Trichostatin A (TSA). Seven out of the eight TSA-treated fragments died during stages 1–4 of the regeneration process; while one fragment survived, it had only reached stage 4 by the 18-day mark (Fig. S3). Control fragments (n = 10) showed a better regeneration response with four fragments fully regenerating to stage 5 (fully functional adult). Three fragments were in the process of regenerating but looked unhealthy by day 18 and three fragments had died (Fig. S3). This could be partly due to the addition of ethanol to the sea water: ethanol was used to dilute the TSA; therefore, the same volume of ethanol was added to the controls. The results of the TSA experiment further supported the conclusions of the VPA-treatment experiments (Fig. 4), showing that HDAC is essential for WBR to successfully occur in *B. leachii*, as in both experiments no fragments were able to regenerate a fully functional adult.

### VPA treatment alters regeneration gene expression

To analyse the effect of reducing HDAC activity on gene expression levels, the expression of key regeneration genes (Zondag et al., 2016) was analysed in *B. leachii* fragments exposed to VPA in comparison to control fragments during regeneration (Fig. 6A). These genes were chosen because, either we had previously identified that they were differentially expressed early on during regeneration (Zondag et al., 2016), or they have been linked to WBR in *B. leachii* elsewhere (Rinkevich et al., 2008; Rinkevich et al., 2010). RNA was extracted between stages 1 and 2 (~15 - 24 h) of WBR, as this was the stage that all VPA fragments were still healthy (Fig. 4).

**Figure 6:**
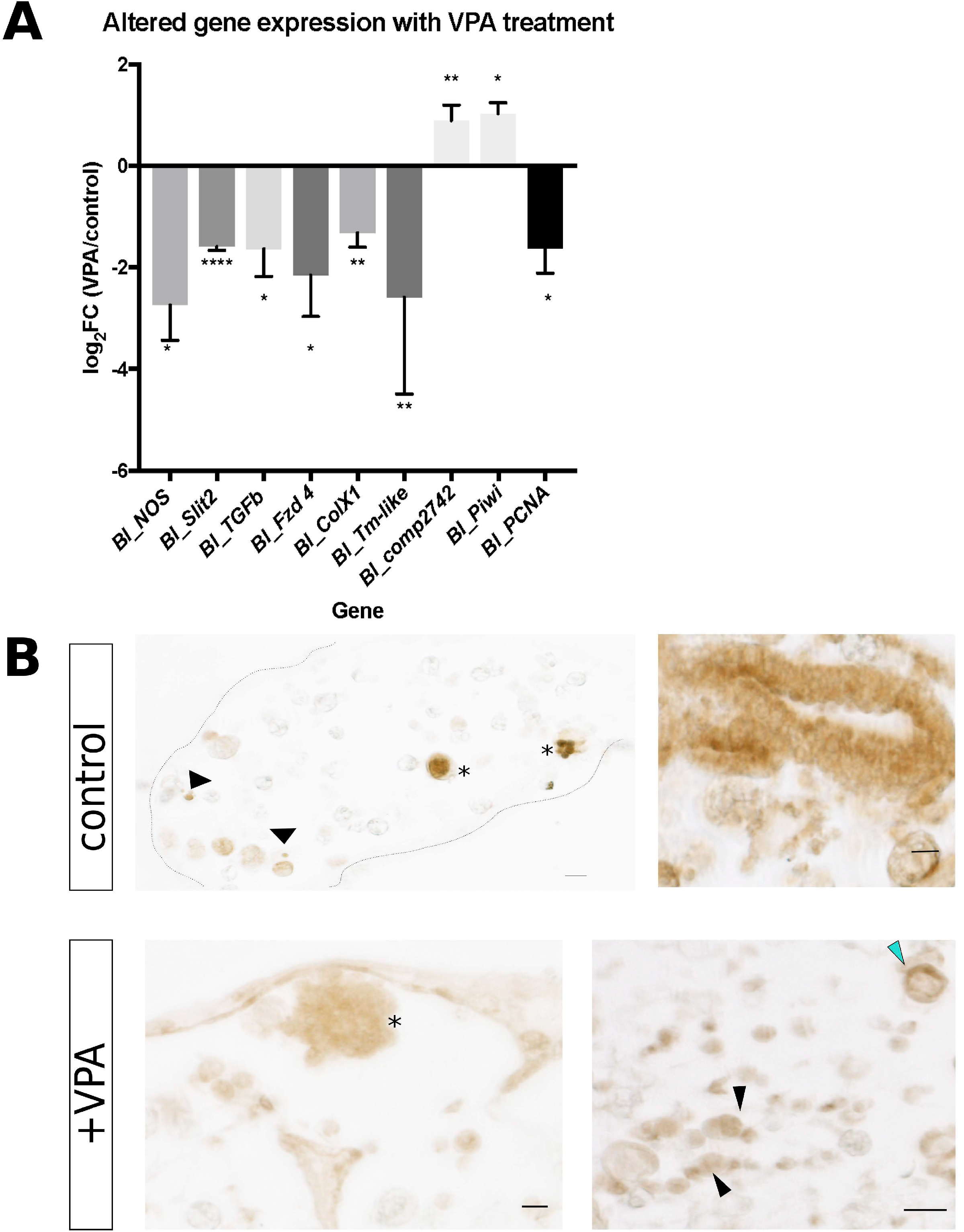
Altered gene and PCNA expression upon inhibition of HDACI/II activity. RT-qPCR and immunostaining for PCNA revealed molecular changes upon HDACi. **A**. Fold change in gene expression of *Bl_Fzd, Bl_NOS, Bl_Slit2, Bl_TGF-ß, Bl_Piwi, Bl_PCNA Bl_Tm-like* and *Bl_ColX1* in VPA-treated tissue undergoing WBR compared to matched controls. VPA-treated and control tissue were collected between 15 - 24 h after removal of all zooids. Data shows mean ± SE, unpaired two-tailed t-test P < 0.05 *, P < 0.005 **, P< 0.0001 ****. **B**. PCNA staining of control and VPA-treated fragments, collected at 48 h post zooid removal. Strong staining was detected in cell aggregates (asterisks) and small individual cells that resemble haemyoblast cells (arrowheads). Cells of the regenerating bud also stained for strongly for PCNA (right panel). VPA-treated fragments (+VPA) showed weaker staining for PCNA overall, even within large cell aggregates (asterisk) and smaller differentiating cells (black arrowheads). Some positive staining was also observed for macrophage-like cells (blue arrowhead). Scale bar is 20 μm.

*Bl_ColX1* (*comp17500*) is predicted to encode a fibrillar collagen isoform X1 protein (ColX1). Previously, we found that gene expression of *Bl_ColX1* significantly declined during the first 24 h of WBR (Padj < 0.05, (Zondag et al., 2016)). Treatment with VPA resulted in a further reduction of *Bl_ColX1* gene expression compared to matched control samples (Fig. 6A; P = 0.01, unpaired two tailed t-test). This suggests that reduced HDAC activity leads to a further suppression of *Bl_ColX1* gene expression.

*Bl_Tm-like* (*comp2742*) is predicted to encode a metalloproteinase (MMP) with thrombospondin motifs (Zondag et al., 2016). MMPs are required for remodelling of the extracellular matrix (ECM) during regeneration in vertebrates (Bai et al., 2005; Vinarsky et al., 2005). *Bl_Tm-like* transcript expression increased more than 3-fold following induction of WBR (Fig. 6A; Padj < 0.05, (Zondag et al., 2016)). Fragments regenerating in the presence of VPA further increased gene expression by almost 2-fold compared to control regenerating fragments (P = 0.03; Fig. 6A).

The enzyme nitric oxide synthase (NOS) produces nitric oxide, a small molecular regulator of many biological processes, including tissue regeneration (Jaszczak et al., 2015; Rai et al., 1998). Previously, we found *Bl_NOS* gene expression increased ~3 fold during early WBR (comp16908, Padj < 0.05; (Zondag et al., 2016)). HDAC inhibition resulted in a decrease of *Bl_NOS* gene expression by 2.7-fold (P = 0.02), compared to control fragments (Fig. 6A).

*Bl_TGF-*□ gene expression is upregulated significantly 3-fold during early WBR (Fig. S4; stages 0 – 2 (Zondag et al., 2016)). In contrast, inhibition of HDAC activity by VPA significantly reduced expression of *Bl_TGF-*□ (P = 0.03; Fig. 6A). The TGF□ pathway is required for regeneration in both vertebrates and invertebrates (Chablais and Jazwinska, 2012; Chen and Xu, 2017; Karkampouna et al., 2012; Petersen et al., 2015).

Components of both canonical and non-canonical Wnt signalling were down- and up-regulated respectively, between early and late stages of WBR (Zondag et al., 2016). Expression of the Wnt receptor *Bl_Frizzed-4* (*Bl_Fzd4*) declines during early WBR and then significantly increases between stages 3 and 5 (Fig. S4, Padj = 0.02). HDACi resulted in a ~2-fold reduction of *Bl_Fzd4* gene expression (Fig. 6A, P = 0.04), suggesting that HDAC activity is required for *Bl_Fzd4* gene expression.

*Slit* genes encode proteins important for axon guidance during development but are also reactivated during axon regeneration in planarians (Cebria et al., 2007). *B. leachii Slit2* mRNA is expressed by circulatory cells within early stage regeneration niches (Rinkevich et al., 2008) and its expression increases between early and mid-WBR (Fig. S4, Padj < 0.05). Regeneration in the presence of VPA reduced *Bl_Slit2* gene expression by 2.2-fold (P < 0.001; Fig. 6A).

Proliferating cell antigen (PCNA) has been used as a marker of cell proliferation during *B. leachii* WBR (Rinkevich et al., 2007). Circulating haemocytes stain with PCNA during the earliest phase of regeneration, and, at two days post regeneration induction, PCNA staining was observed in cell aggregates within regeneration niches (Rinkevich et al., 2007), indicating that cell proliferation is occurring during these initial regeneration stages. As HDAC activity promotes cell cycle progression and cell proliferation in some cell types and can directly interact with PCNA (Bhaskara et al., 2013; Glozak and Seto, 2007; Milutinovic et al., 2002), we determined if *PCNA* mRNA expression was affected by treatment with VPA. *Bl_PCNA* mRNA levels decreased 1.6-fold with VPA treatment (Fig. 6A, P = 0.03), suggesting that HDAC activity was needed for correct *PCNA* gene expression during WBR. To further investigate this, we stained regenerating fragments with antibodies against PCNA (Fig. 6B). Control fragments showed strong staining for PCNA in cell aggregates and regenerating buds (Fig. 6B). However, PCNA staining of VPA-treated fragments showed weaker signal, including within cell aggregates (Fig. 6B), suggesting reduced levels of PCNA protein.

*Piwi* genes encode RNA-binding proteins, expressed in both the germ line and adult pluripotent stem cells (van Wolfswinkel, 2014). The *Bl_Piwi* gene plays a key role in WBR; its expression increases during WBR, initially in cells that line the blood vessel epithelium and later in cells through the vessel lumen (Rinkevich et al., 2010). *Bl_Piwi* gene knockdown by siRNA arrests regeneration, and these fragments lack regeneration buds and cell aggregates (Rinkevich et al., 2010). HDAC inhibition leads to increased expression of *Bl_Piwi*, in comparison to control fragments (P = 0.008; Fig. 6A), suggesting that inhibition of HDAC leads to dysregulation of *Bl_Piwi* gene expression. In situ hybridisation for Bl_Piwi transcript showed that in control fragments, it was highly expressed in haemoblasts and larger cell aggregates (Fig. 7A and B). Following exposure of WBR to VPA, *Bl_Piwi* expression was now found in cells throughout the haemolymph (Fig. 7C), including immune cell types (Fig. 7D).

**Figure 7:**
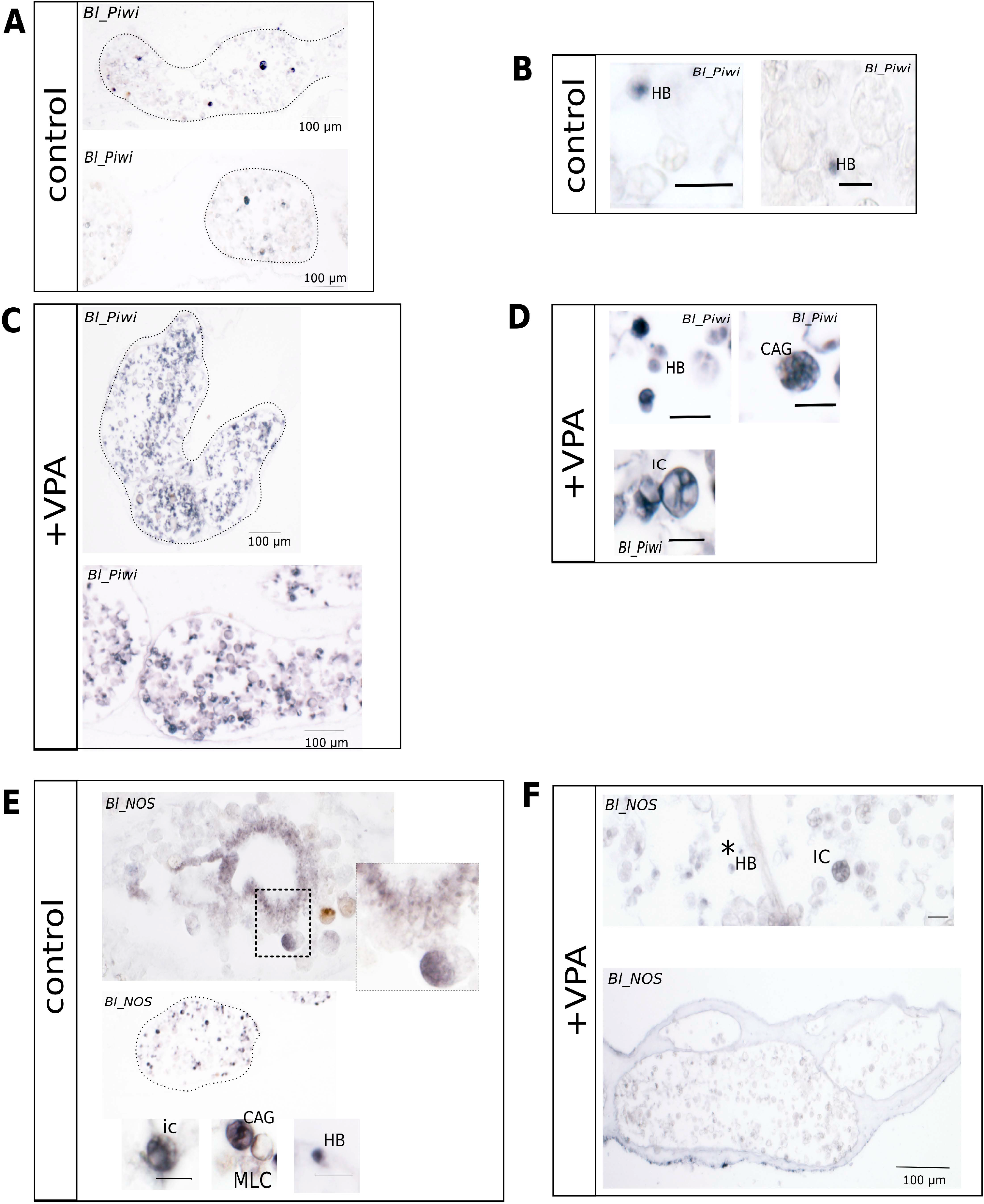
*Bl_Piwi* and *Bl_NOS* gene expression following HDACi. *In situ* hybridisation for *Bl_Piwi* and *Bl_NOS* mRNA during WBR in control and VPA-treated fragments. **A**. Control WBR *Bl_Piwi* expression was detected in cells located in some of the condensed ampullae. **B**. Small cells expressing *Bl_Piwi* mRNA, surrounded by much larger haemocytes that do not stain for *Piwi*. These smaller cells are putative stem-like cells (Rinkevich et al., 2010). **C**. *Bl_Piwi* expression is detected in cells throughout the ampullae following VPA-treatment during WBR. **D**. Cells expressing *Bl_Piwi* include the smaller haemyoblast cells, cell aggregates and larger vacuole containing immunocytes, with a morphologies similar to macrophage and morula cell types. **E**. *Bl_NOS* mRNA was detected in haemoblasts and morula cells and cell aggregates as well as the epithelium of the regenerating bud (enlargement shown). **F**. VPA treatment reduced overall staining for *Bl_NOS* mRNA, with some gene expression still detected in haemyoblast (asterisks) and morula cells. Abbreviations: haemoblast (HB), morula cell (MC), cell aggregate (CAG), macrophage-like cell (MLC), immunocyte (IC). Scale bars are 10 μm, unless otherwise shown. Identification of cell types was based on (Blanchoud et al., 2017).

In comparison, *Bl_NOS* gene expression, which decreased with HDACi (Fig. 6A), was more difficult to detect by *in situ* hybridisation following VPA treatment (Fig. 7F). In control fragments, *Bl_NOS* mRNA was detected in immunocytes, cell aggregates and haemoblasts, and the epithelium of regeneration buds (Fig. 7E). In the VPA-treated fragments, *Bl_NOS* was only just detectable above background, enriched in haemoblasts and some cell aggregates (Fig. 7F).

### *HDAC2, HDAC9* and *Kat8* genes are expressed in most cell types during WBR

As WBR is halted mid-way through by HDACi, resulting in dysregulation of gene expression, we determined the corresponding spatial expression of *Bl_HDAC2* and *Bl_Kat8* mRNA at mid-WBR (stage 3; Fig. 8). *Bl_HDAC2* mRNA was expressed by multiple immune-like cells in *B. leachii* circulation at stage 3, including morula cells (Fig. 8A). Only weak staining was found in the regeneration bud, located at the posterior end of the bud. Cell aggregates located near the bud also did not express *Bl_HDAC2* in comparison to the morula cells (an immunocyte cell lineage) (Fig. 8A).

*Bl_HDAC9* mRNA expression was detected in most immunocytes, transport and pigment cells (Fig. 78B). It is also expressed by cells that appeared to be differentiating cells, but not the mature haemoblast cells themselves (Fig. 8B). The differentiating cells (DCs) often have a larger cytoplasm than the haemoblast cells, and are thought to be transitory cells (Blanchoud et al., 2017; Cima et al., 2002). *Bl_HDAC9* mRNA was found in epithelial cells, located in the posterior region of the regeneration bud and some nearby cell aggregates (Fig. 8B). *Bl_Kat8* was also expressed broadly in the immunocytes, differentiating cells and haemoblasts (Fig. 8C). The regeneration bud also showed higher expression, in both the posterior epithelium and some cell aggregates surrounding the developing bud (Fig. 8C).

Expression of the *Bl_Piwi* expanded to immunocytes upon HDACi (Fig. 7A-D), correlating with inhibition of HDACI/II type enzyme which are strongly expressed within these cell types (Fig. 8A-B). *Bl_NOS* gene expression declined upon VPA treatment, *Bl_NOS* mRNA was detected in multiple immunocytes as well as the regeneration bud. Failure to form the regeneration bud and/or reduced expression of *Bl_NOS* in cells that typically express HDACI/II enzymes could be responsible for the reduction of *Bl_NOS* gene expression.

## Discussion

### Identification and expression of HDAC and HAT genes during WBR

Examination of the *B. leachii* genome, alongside additional ascidian genomes revealed that each ascidian genome had two Class I, two Class II and a single Class IV *HDAC* gene. This is also true for other invertebrates such as *Drosophila*, providing further evidence that the metazoan ancestor genome contained two Class I, two Class II and a single Class IV *HDAC* gene, and that the increase in vertebrate *HDAC* gene numbers is due to gene duplication (Gregoretti et al., 2004). Each vertebrate paralogue have very similar protein sequences (particularly HDAC1 and 2), and are often shown to function redundantly in development (Jurkin et al., 2011). The ascidian genomes examined also contained a single representative for each HAT/KAT subclass.

HDAC and HAT function in regulating chromatin structural changes and DNA accessibility to globally control gene expression from the genome. Two important aspects that restrict function of these proteins are both their overall availability (expression level), and their enzymatic activity (Haberland et al., 2009; Legube and Trouche, 2003). As such, we focused on *HAT* and *HDAC* genes highly expressed during WBR, assuming this indicates that they have important roles in this process. While RNA analysis does not factor in post-translational changes such as phosphorylation, known to alter enzymatic activity (Legube and Trouche, 2003), any dynamic changes to mRNA expression levels is likely to have a functional consequence for genome expression during regeneration. The differences in expression levels between family members is likely due to divergent cellular functions. Class I HDAC proteins are mostly confined to the cell nucleus (Delcuve et al., 2012; Seto and Yoshida, 2014), and are important for cell proliferation, regulation and differentiation (Reichert et al., 2012), whereas class II HDAC proteins shuttle between nucleus and cytoplasm and are thought to have more tissue specific functions (Delcuve et al., 2012; Seto and Yoshida, 2014). *MYST/KAT* genes are required for stem cells and developmental processes (Sapountzi and Cote, 2011). HAT1 (also known as KAT1) enzymes are associated with DNA replication and repair (Agudelo Garcia et al., 2017; Nagarajan et al., 2013) and are essential for the acetylation of newly synthesised histone H4 and H3 proteins (Nagarajan et al., 2013). The p300 has important roles in cell proliferation, and, together with CREB binding protein (CBP), promotes the acetylation of histone 3 lysine 27 (H3K27) and histone 3 lysine 18 (H3K18) (Bedford et al., 2010; Ogryzko et al., 1996).

The gene expression of the *B. leachii* class II HDAC, *Bl_HDAC2*, peaked at stage 3 (Fig. 3). During stage 3 regeneration niches appear containing epithelial buds, that later form a new zooid. This is also the stage when HDACi stalls regeneration (Fig. 4). *Bl_HDAC2* expression in the circulating haemocytes, restricted to immunocytes, morula cells, with weaker expression within the regeneration bud (Fig. 8). Notably, *B. leachii HDAC* genes were not expressed in all epithelial cells of the regenerating bud, but were enriched within cells located at the posterior end of the bud, across from the developing endostyle (Fig. 8). HDAC activity is required for axis patterning and cell lineage commitment in other animals (Brunmeir et al., 2009; Carneiro et al., 2011; Lv et al., 2014), suggesting *B. leachii* HDAC genes may function in the development of specific tissue layers. This may additionally relate to the observation in HDACi histology sections, where HDACi fragments with a bud present after a week exhibit one end poorly formed. However, it is not possible to distinguish whether one end had either formed and the bud had halted development, or was in the process of degeneration (Fig. 5). HDAC 1/2 complexes have important roles in differentiation of certain cells types and determining the fate of ESCs (Dovey et al., 2010; Turgeon et al., 2013; Ye et al., 2009). It is possible that HDACi, resulted in a change of cell lineage commitment, promoting one cell fate over another, resulting in failure of the regeneration bud to continue development.

**Figure 8.**
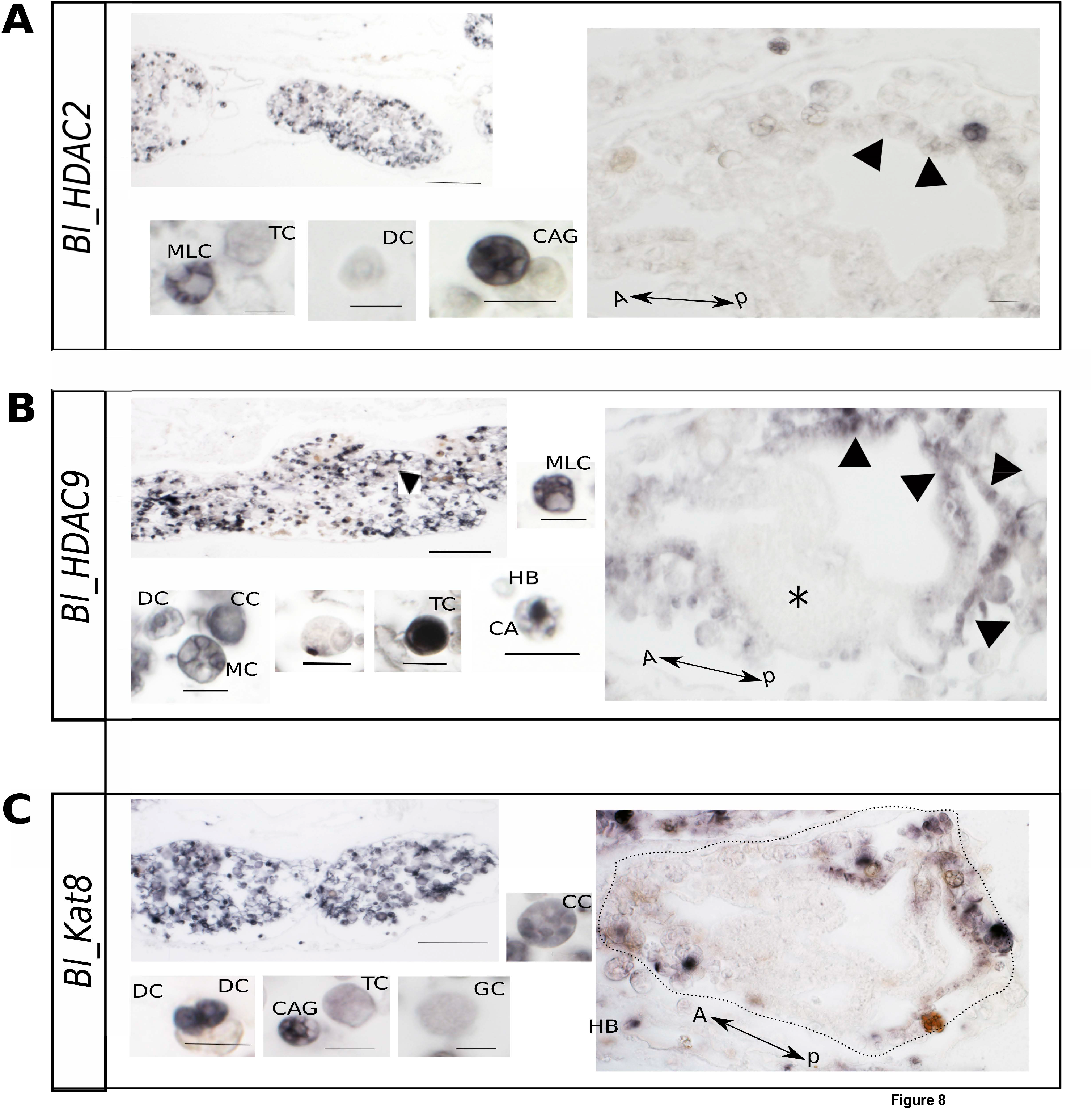
Expression of *Bl_HDAC2, Bl_HDAC9* and *Bl_Kat8* in control WBR fragments. *In situ* hybridisation was used to detect expression of target transcripts in sectioned WBR fragments collected two days after removal of all zooids. **A**. *Bl_HDAC2* mRNA was detected throughout the ampullae, in immunocytes, cell aggregates but not differentiating cells. Gene expression was also detected weakly in the dorsal regions of the regeneration bud. **B**. *Bl_HDAC9* mRNA was strongly expressed by most circulating cell populations including immunocytes and transport cells (CA). It was also expressed strongly in the cells of the regenerating bud, mainly at the dorsal end (arrow heads) and absent from cells aggregating in the middle (asterisk). **C**. Staining for *Bl_Kat8* mRNA was found in multiple circulating haemocytes including transport cells (TC), small differentiating cells and cell aggregates (CAG). It was expressed by haemocytes near the regeneration bud and by cells at the dorsal end of the bud (arrowheads). Abbreviations: haemoblast (HB), morula cell (MC), cell aggregate (CAG), immunocyte (IC), anterior (A), dorsal (V), transport cell (TC), differentiating cell (DC). Scale bars are 10 μm, unless otherwise shown.

**Figure 9.**
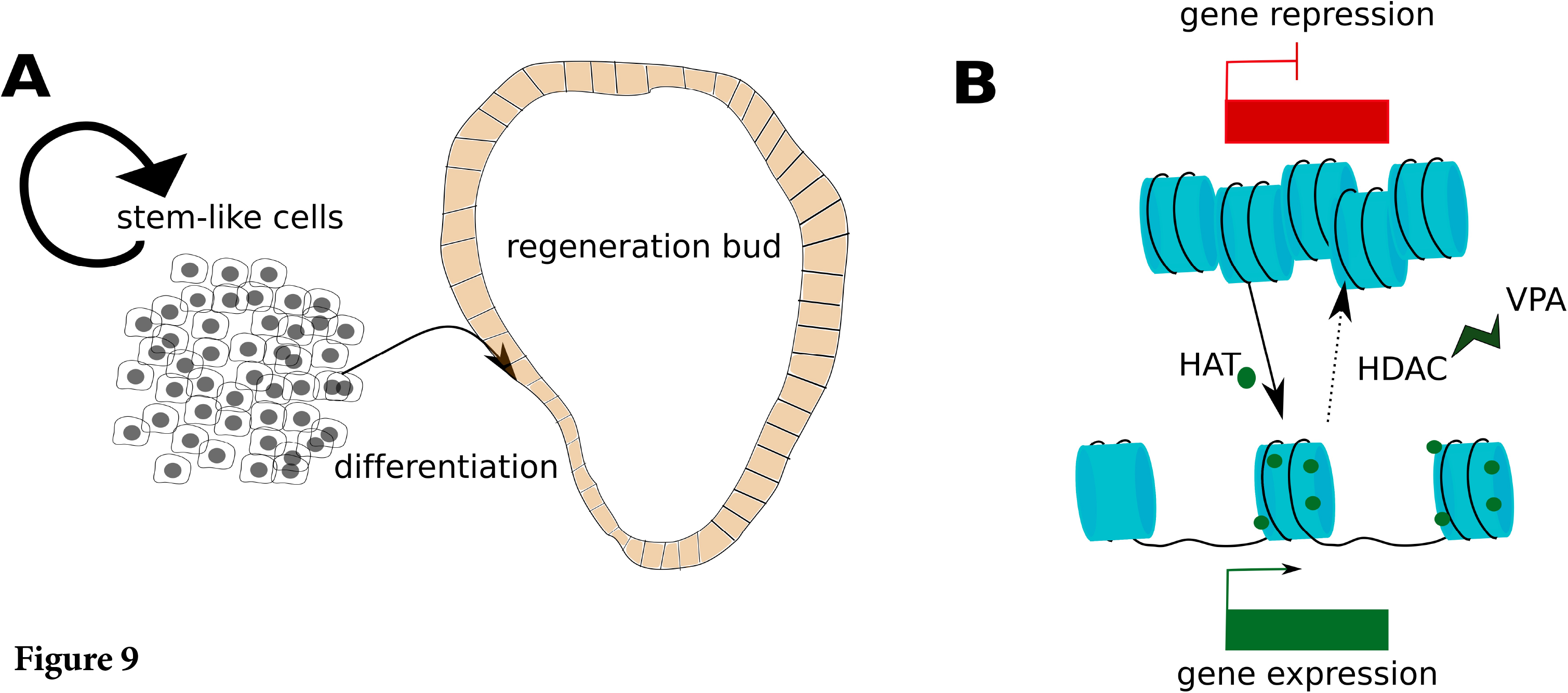
Model of HDAC inhibition. Schema depicting a model for *B. leachii* WBR and histone acetylation by HDAC/HAT enzymes. **A**. Hypothesis for *B. leachii* regeneration based on current and earlier studies (Rinkevich et al., 1995; Rinkevich et al., 2010). WBR is initiated with an increase in stem-like cells (Piwi positive cells) (Rinkevich et al., 2010). These form a cell aggregate inside a regeneration niche; an area where several ampullae have fused. Around stage 2–3, ~ 24–48 h post-zooid removal, cells exit this aggregate and form an epithelial regeneration bud before differentiating into specialized cell types found in a mature zooid. **B**. HAT and HDAC enzymes regulate gene expression from the *B. leachii* genome, compact (heterochromatin) and open forms of chromatin (euchromatin). HDAC inhibition (by VPA) increased acetylation of nuclear proteins, leading to halting of WBR and altered gene expression including those genes required for differentiation.

KAT8 (MYST1/MOF) proteins have multiple cellular roles including controlling cell cycle progression (Thomas et al., 2008), pluripotency and regulating expression of core ESC genes (Li et al., 2012). *Bl_KAT8* gene expression peaks at stage 3 (Fig. 3), and was detected in immunocytes, hemoblasts and differentiating cells (Fig. 8).

The *p300* gene functions to regulate cell proliferation and differentiation in many animals, and also acts as a transcriptional co-activator with many transcription factor proteins (Bedford et al., 2010; Kasper et al., 2002). HAT proteins such as p300 promote transcriptional activation by creating binding sites for chromatin remodelling complexes to bind and open up the DNA (Bedford et al., 2010). *KAT7* genes (also known as *Myst2/Hbo1*) regulate the expression of genes required for embryonic development (Kueh et al., 2011) and mouse embryonic stem cells (Pardo et al., 2017). KAT7, in combination with partner proteins, targets specific lysine residues on H3/H4 histone proteins for acetylation (Pardo et al., 2017; Sapountzi and Cote, 2011).

While the focus here is regeneration, it is likely that HDAC activity is also required for asexual reproduction in *B. leachii*. Later stages of WBR are likely to be similar to asexual reproduction, particularly following formation of a regeneration bud, which is morphologically similar to the blastogenesis budlet (Kurn et al., 2011). GCN5 (also known as KAT2a) activity is required for blastogenesis in *P. misakiensis* but not zooid regeneration (Shibuya et al., 2015). We also note that *KAT2* expression is lower compared to other *HAT/KAT* genes, suggesting it may not have a key role in colonial ascidian regeneration. As there is functional redundancy between HAT/KAT proteins (Nugent et al., 2010), we predict that use of a broader HAT inhibitor would have also halted regeneration in *P. misakinensis*. However, in the future it would be of interest to use specific knockdown of each *KAT* gene to determine if they have specific roles in asexual reproduction and regeneration.

### Changes to WBR gene expression resulting from HDACi

HDACi altered the expression of genes typically up- or down-regulated during WBR, indicating changes to gene expression is likely to have played a role in failure of regeneration failure observed with VPA and TSA treatment. While HDAC activity is more commonly associated with progressing chromatin into a more compressed heterochromatin structure, it can also cause an increase in gene expression due to suppression of repressor gene expression (Glaser et al., 2003). This suggests that the correct implementation of WBR-associated gene expression patterns requires HDAC function and protein deacetylation.

Modification of the ECM is important for facilitating regeneration (Bonnans et al., 2014). HDACi did alter expression of these genes, their expression altered in a similar direction (up or down) to that of normal WBR. Additionally, given that HDACi did not halt condensation and reorganisation of the ampullae (Fig. 3 and 4), this further suggests that HDAC activity and protein deacetylation does not play an essential role in the reorganisation of the vascular tissue during early WBR.

Cell proliferation is an important factor in regeneration, providing a source of cells for the generation of replacement tissues (Sanchez Alvarado and Tsonis, 2006). In our study, HDACi decreased PCNA levels as determined by both RT-qPCR and IHC (Fig. 6). Previous studies investigating mouse liver and kidney regeneration also showed that HDAC function is required for increasing PCNA levels (Tang et al., 2014; Wang et al., 2008), indicating that this may not just be a feature of WBR but many types of regenerative processes across different phyla.

*Bl_Slit2* gene expression typically increased between stages 0 to 3 of WBR. HDACi significantly reduced *Bl_Slit2* mRNA levels, indicating it HDAC activity is required for up-regulation of *Bl_Slit2* gene expression. Slit2, in addition to roles in axon guidance, also functions in the formation of vascular structures and in vascular repair following injury (Yuen and Robinson, 2013). *Slit2* gene expression is directly regulated by HDAC5 (a class II HDAC) in mammalian endothelial cells (Urbich et al., 2009). This gene may function in regulating the vascular rearrangements that occur during *B. leachii* WBR.

*Bl_NOS* gene expression also reduced with HDACi treatment. NOS is essential for the generation of NO, which has important roles in pluripotency, mesoendoderm specification, and in immune responses (Beltran-Povea et al., 2015; Cencioni et al., 2018; Lee et al., 2017). *Bl_NOS* mRNA is expressed in multiple cell types including hemoblasts and the regeneration niche. Thus, its expression pattern during WBR suggests that nitric oxide also has important roles in these biological processes within *B. leachii* colonies. The reduction of NOS expression by HDACi could be due to loss of regeneration niches (that expression NOS) and/or a reduction of certain NOS expression cell types following HDACi treatment.

### Hypothesis and model for HDACI/II function in *B. leachii* WBR

During *B. leachii* regeneration, a new bud forms located within a regeneration niche, beside an aggregate of stem-like/hemoblast cells that express the pluripotency gene *Piwi* (Fig. 7A). The acetylation and de-acetylation of nuclear proteins is important for correct gene expression during development and regeneration (Huang et al., 2014; Taylor and Beck, 2012; Tseng et al., 2011). Thus, loss of HDAC activity halts regeneration between stages 3 and 4, a critical period in which changes to gene expression is required to induce cells to undergo differentiation and drive tissue patterning into a new zooid. Previous examination of differential gene expression found the expression of over 500 transcripts declined between stages 3 and 4 ((Zondag et al., 2016); Fig. S5), compared to increased expression of only 39 genes; this potentially could be a result of histone de-acetylation and the expression of repressor proteins. Future studies will include RNA-sequencing (following HDACi), alongside chromatin immunopreciptation experiments, for a more global approach to identifying targets of histone acetylation during WBR.

Given the importance of histone deacetylation in gene regulation and differentiation, it may be a conserved method utilised by many animals to regulate changes to global gene expression from the genome during regeneration. Reduced HDAC activity also prevents formation of the blastema in cnidarian *Hydractinia* head regeneration by both HDACi and HDAC2 knockdown (Flici and Frank, 2018; Flici et al., 2017). In vertebrates, HDACi prevents hindlimb muscle regeneration in mice (Spallotta et al., 2013), and tail and limb regeneration in *Xenopus* (Taylor and Beck, 2012; Tseng et al., 2011). Chemical inhibition of HDAC function in zebrafish does not interfere with the initial stages of fin regeneration (healing and blastema formation), but causes later defects, preventing differentiation of cells and reducing tail outgrowth (Pfefferli et al., 2014). We also found that HDACi did not block the initial stages of regeneration, instead stopping regeneration after the ampullae had condensed to form the regeneration niches, at the bud formation and differentiation stage. Therefore, while all these animals all use different modes of regeneration, like blastema formation, cell dedifferentiation and stem cells to repair and replace lost tissue, they all require HDAC function for successful regeneration.

## Conclusions

We observed dynamic changes in the expression patterns of the various epigenetic modifiers observed during WBR in *B. leachii*. Well-coordinated regulation of these epigenetic enzymes allows the expression of the appropriate sets of genes during the course of regeneration. Inhibition of HDAC activity caused a dysregulation in gene expression and ultimately led to a failure of the vascular tissue undergoing cellular differentiation during WBR. During the first 24 hours of WBR existing tissue is reorganised including fusion of some ampullae, followed by formation of a ‘regeneration niche’ seen in the vascular fragments at stage 3. This ‘opaque niche’ becomes pigmented as cells differentiate to form the new adult, however this process did not occur in the majority of fragments exposed to VPA. We therefore predict that HDAC is needed to allow the putative ‘stem’ cells to undergo differentiation and/or to proliferate. Although we only focused on the factors with the highest expression, other classes of HDAC enzymes could play a role in a more restricted manner, for instance in a subset of specific cells. Future work using the *B. leachii* genome (Blanchoud et al., 2018) and mapping of epigenetic marks will aid our understanding of how these epigenetic modifiers influence global gene expression.

## Methods

### Identification *B. leachii* orthologues and phylogenetic analysis

*B. leachii* orthologues of HAT, HDAC and histone proteins were identified using a TBLASTN 2.2.26+ against the entire *B. leachii* transcriptome using conserved protein domain sequences (File S1). Contig identification was additionally confirmed by reciprocal Blast using SMARTBLAST (https://blast.ncbi.nlm.nih.gov/smartblast/). Conserved protein domains used for identification of these domains in HAT/KAT proteins are listed in Supplementary Table 2.

To assign subgroups to *B. leachii* HAT and HDAC orthologues, phylogenetic analysis was carried out with full length sequences. Amino acid sequences were aligned with ClustalX (Jeanmougin et al., 1998), followed by Bayesian analysis using MrBayes (Ronquist et al., 2012) for 10,000 generations under a mixed model for HDAC proteins, and a fixed model (Jones) for HAT, both with a burin of 250. For expression analysis, RNA-seq count data (Zondag et al., 2016) was normalised to library size to determine expression changes across regeneration stages (Supplementary Data File 2).

### *B. leachii* husbandry and regeneration

*B. leachii* colonies were collected from Otago Harbour (latitude 45.87°S, longitude 170.53°E) in New Zealand and cultured as described previously (Zondag et al., 2016).

Regeneration of *B. leachii* zooids from vascular fragments was carried out as previously published (Zondag et al., 2016). All zooids and buds from the marginal ampullae were removed using a fine tungsten needle. The vessel fragments attached to the slides were returned to the aerated saltwater tanks and monitored under a Leica M205 FA stereomicroscope. Imaging was performed using the Leica DFC490 digital camera. Regenerating fragments were then removed from the glass slides for protein or RNA isolation.

### *B. leachii* staging

Staging of regeneration stages was carried out as per Zondag *et al*, (2016); images are shown in Supplementary Figure S1. Briefly Stage A: *B. leachii* colony. Stage 0: Marginal ampullae at 0 h directly after dissection from the zooids has taken place. Stage 1: New vascular connections formed between ampullae, creating the beginning of a new circulatory system, but terminal ampullae are still in initial cone-shape form. Stage 2: Marginal ampullae started to reshape and condense together and/or a compact network of blood vessels within the tunic matrix has formed. Stage 3: Remaining ampullae have completely condensed. Stage 4: Formation of small transparent vesicle in the middle of the condensed blood vessels. This transparent ball continues to expand in size and eventually gains pigment before forming the new zooid. Stage 5: A fully functioning adult zooid capable of filter feeding.

### Chemical inhibition of HDAC activity

VPA was diluted in seawater to give a final concentration of 1mM. Dissected *B. leachii* colonies (leaving only vascular tissue) were submerged in aerated containers filled with 900 mL salt water containing either VPA or seawater only in the control. Seawater and VPA was replaced every other day. WBR was monitored for 18 days.

TSA was diluted in ethanol (EtOH) to a stock concentration of 4 mM. *B. leachii* colonies were dissected (leaving only vascular tissue) and then submerged in aerated containers filled with filtered salt water containing either TSA at a final concentration of 50nM or for the controls (the same volume (10 μl) EtOH without TSA). Seawater and TSA/EtOH was replaced every other day, WBR was monitored for 18 days.

### Regeneration score

WBR in *B. leachii* vascular fragments was scored after 18 days post regeneration induction through dissection of adults. Regeneration scoring was used as a way of quantifying regeneration success. The score reflected the regeneration stage the tunic tissue had reached after 18 days, and if either the vascular fragments either looked healthy or were in the processes of dying or were dead. The varied regeneration abilities were classed between 0–10. For example, if the vascular fragment was healthy and had regenerated to an adult, a regeneration score (RS) of 10 was given. However, if the tissue had died in the first stages (stages 1–2) of WBR it was classed as a 0 (failed regeneration). Each of the regenerating tissue fragments was assigned a RS that was used to determine the effect of VPA on regeneration. A Mann-Whitney test was carried out comparing the VPA exposed fragments to the controls.

### RT-qPCR gene expression analysis

RT-quantitative PCR (RT-qPCR) was carried out using total RNA extracted from VPA exposed vascular tunic fragments or a parallel-run controls. Both tissues were collected between 15 - 24 h post WBR induction. RNA extractions and cDNA synthesis (500 ng of total RNA) were performed as described previously (Zondag et al., 2016). All samples were assayed in triplicate with the SYBR Select Master Mix (ThermoFisher). The RT-qPCR reaction protocol consisted of 50°C for 2 min, 96 °C for 2 min, (40 cycles of 96 °C for 15 s, 55–60 °C for 30 s and 72 °C for 1 min), followed by a dissociation curve program. RT-qPCR data was analysed with the ΔCt (change in cycle threshold) method using 2^-(ΔCt)^ to work out relative RNA expression to the reference genes, *Rpl27* and *Rsp29*. Gene expression fold change was calculated by determining the Log_2_(VPA-treated/control) using relative gene expression levels. A list of RT-qPCR oligonucleotide sequences is provided in Supplementary Table 3.

### Measurement of nuclear acetyl-proteins

The Plant Nuclei Isolation/Extraction Kit (CelLyticPN, Sigma-Aldrich) was used to carry out the nuclear protein extractions. Tissues used for extraction were matched VPA-treated and control tissue undergoing WBR and both were collected at 15 h post adult dissection. Antibodies used for the dot blot were: primary antibody goat polyclonal anti-pan-Acetyl (Santa Cruz) (sc-8649) and secondary antibody Donkey anti-goat-800 (LI-COR Biosciences; 925–32214).

Protein extracts were diluted in 1x SDS buffer (□-mercaptoethanol (0.1%), bromophenol blue (0.0005%), glycerol (10%), SDS (2%) in 63 mM TrisCl pH 6.8) to produce five, 2-fold serial dilutions. One □1 of diluted protein was placed onto a nitrocellulose membrane and left to dry for 30 min. Membranes were then soaked in REVERT (LI-COR Biosciences) stain for 5 min and washed twice with REVERT wash buffer prior to scanning at 700nm. The membrane was then blocked in Odyssey blocking buffer for 30 min, followed by addition of the anti-pan Acetyl antibody diluted to 1 in 1000 in blocking buffer. Following an overnight incubation at 4°C with rocking, the membrane was washed three times with PBTw (phosphate-buffered saline with Tween-20 (0.5%)) for 10 min each. The secondary antibody, anti-goat-800 (1 in 5000) was added and the membrane left to incubate for 1 h at room temperature. The membrane was washed three times with PBTw, then a single wash with PBS, left to dry membrane before scanning at 800nm. REVERT total protein stain (LI-COR Biosciences) total protein stain used for normalization of protein levels. Membranes were scanned using the Odyssey Imaging System. For protein quantification, signal intensity values were used to normalize anti-pan-Acetyl antibody staining to total protein signal using ImageStudioLite application (LI-COR Biosciences).

### Histology

Control and VPA-treated fragments were fixed in 4% PFA for 1 h at room temperature. They were then removed from the glass slides and washed once in PBS before dehydration and embedding in paraffin, orientated for sagittal sections. Paraffin blocks were cut into 5-μm sections by the Histology unit (University of Otago). Sections were cut at 5 □m and stained with hematoxylin and eosin (H&E stain) using standard methods. Stained sections were imaged with a Olympus AX70 light microscope using 20x and 40x objectives.

### PCNA staining

*B. leachii* sections were deparaffinized and rehydrated through an ethanol:water dilution series. Antigen retrieval was carried out with slides immersed in 10 mM sodium citrate buffer (pH 6.0) and microwaved on high for 30 min. Slides were then rinsed with ddH_2_0 several times before continuing onto the immunohistochemistry protocol. Slides were then washed several times with PBTx (PBS plus 0.025% Triton X-100), for 5 min each. Non-specific binding sites were blocked with PBS containing 1% BSA for 2 h. PCNA primary mouse monoclonal conjugated to horse radish peroxidase (PC10, Santa Cruz (sc-56-HRP)) was diluted to a final concentration of 1 in 500 in blocking buffer before addition to the slides. Following an overnight incubation at 4°C, slides were washed three times with PBTx for 5 min each. Staining was carried out using the Invitrogen DAB plus kit according to the manufacturer’s instructions. The color reaction was stopped by washing the slides with ddH_2_0 and then they were mounted for imaging with an Olympus AX70 microscope using a 40x objective.

### *In situ* hybridisation

Primers were designed to amply specific gene fragments from *B. leachii* cDNA (Supplementary Table 3). PCR products were then cloned into pCRII-TOPO (Life Technologies) and sequenced to confirm the correct insertion of the product. Dioxygenin (DIG)-labelled sense and anti-sense RNA probes were synthesised using 10x DIG RNA labelling mix and SP6/T7 RNA polymerases (Roche) in *in vitro* transcription reactions.

Different stages of regenerating *B. leachii* fragments were fixed in 4% paraformaldehyde and dehydrated in 70% methanol, embedded in paraffin and sectioned to 5 μm. Hybridisation of probes to tissue sections was performed using (Breitschopf et al., 1992) method for paraffin-embedded tissues with the following changes. Briefly, sections were rehydrated through a xylene-ethanol series into PBS, fixed with 4% PFA and incubated with proteinase K (□□g/ml in PBS) for 10 min, followed by re-fixing with 4% PFA for 10 min. Following three washes with PBS, slides were acetylated and then incubated with hybridisation buffer for 1 h. DIG-labelled probes were added to 200 μl of hybridisation solution (50% formamide, 5x saline sodium citrate buffer (SSC), 2% blocking reagent (Roche), 0.1% Triton X-100, 0.5% CHAPS) and slides were incubated overnight at 60°C.

Following washing with 5x SSC and 0.2x SSC at room temperature, slides were blocked with TNB buffer (50 mM TrisCl pH 7.5, 150 mM NaCl with 10% sheep serum) for 1 h before incubation with anti-DIG conjugated with alkaline phosphatase (Roche), diluted 1 in 2000 in TNB buffer overnight at 4°C. Slides were washed three times in TN buffer (50 mM TrisCl pH 7.5, 150 mM NaCl) and finally with NTM buffer (10 mM TrisCl pH 9.5, 50 mM MgCl_2_, 0.1 M NaCl). The colour reaction solution contained NBT/BCIP in NTM. Once sufficiently stained, slides were washed briefly in PBS plus 1% Triton X-100, fixed with 4% PFA and imaged with an Olympus AX70 Light microscope.

## List of abbreviations

HDAC: Histone deacetylase
HAT: histone acetyl transferase
WBR: Whole body regeneration
CPTH2: (cyclopentylidene [4’-(4-chlorophenyl)thiazol-2-yl) hydrazone)
FPKM: Fragments per kilobase per million
HDACi: HDAC inhibition

## Declarations

### Ethics approval

not applicable

### Consent for publication

not applicable

### Availability of data and material

Data sharing is not applicable to this article as no datasets were generated or analysed during the current study that are not provided with the article.

### Author’s contributions

LZ carried out the inhibitor experiments, transcriptome analysis and some of the RT-qPCR experiments. RC performed the majority of the RT-qPCR and some of the *in situ* hybridisation experiments. MJW carried out the histology and in situ hybridisation experiments. All authors contributed to writing of the manuscript.

## Acknowledgments

We thank James Smith, Lyvianne Decourtye, Jeremy McCallum-Loudeac, Susie Szakats, Michael Meier and Stephanie Workman for feedback on manuscript drafts. LZ was supported by a University of Otago PhD scholarship.

